# The neocortical infrastructure for language involves region-specific patterns of laminar gene expression

**DOI:** 10.1101/2024.01.17.575815

**Authors:** Maggie M.K. Wong, Zhiqiang Sha, Lukas Lütje, Xiang-Zhen Kong, Niels Velthuijs, Sabrina van Heukelum, Wilma D.J. van de Berg, Laura E. Jonkman, Simon E. Fisher, Clyde Francks

## Abstract

The language network of the human brain has core components in the inferior frontal cortex and superior/middle temporal cortex, with left-hemisphere dominance in most people. Functional specialization and interconnectivity of these neocortical regions is likely to be reflected in their molecular and cellular profiles. Excitatory connections between cortical regions arise and innervate according to layer-specific patterns. Here we generated a new gene expression dataset from human postmortem cortical tissue samples from core language network regions, using spatial transcriptomics to discriminate gene expression across cortical layers. Integration of these data with existing single-cell expression data identified 56 genes that showed differences in laminar expression profiles between frontal and temporal language cortex together with upregulation in layer II/III and/or layer V/VI excitatory neurons. Based on data from large-scale genome-wide screening in the population, DNA variants within these 56 genes showed set-level associations with inter-individual variation in structural connectivity between left-hemisphere frontal and temporal language cortex, and with predisposition to dyslexia. The axon guidance genes *SLIT1* and *SLIT2* were consistently implicated. These findings identify region-specific patterns of laminar gene expression as a feature of the brain’s language network.

## Introduction

The human capacity for language relies on a distributed network of brain regions, core to which are the inferior frontal gyrus and superior/middle temporal cortex (1–3). In most people, there is functional dominance of the left-hemisphere regions for language, especially for sentence production (4). A major challenge for biological studies of language is to understand how this regional specialization is supported by distinct molecular and cytoarchitectonic profiles. For example, it is likely that molecules such as transcription factors, neurotransmitter receptors, ion channels, and synaptic adhesion proteins affect neural signaling and inter-areal connectivity, to influence regional functional specialization (5–7).

Postmortem transcriptomic analysis generates a quantitative profile of gene expression values across thousands of active genes within a tissue sample. This approach relies on sampling tissue from defined anatomical regions within a few hours of death, before levels of messenger RNA (mRNA) become too degraded (8–10). Through combining this type of postmortem data with brain maps derived from non-invasive imaging of living individuals, numerous studies have found that patterns of gene expression across the cerebral cortex co-vary with anatomical and functional organization (6, 7, 11–19). Some general principles are that i) cortical transcription profiles exhibit macroscale gradients that correlate with hierarchical specialization, from primary sensorimotor to multimodal association regions, and ii) cortical regions with relatively higher inter-connectivity tend to have somewhat similar gene expression profiles, even when spatially separated.

However, previous analysis that linked postmortem gene expression to connectivity within the human brain’s left-hemisphere language network relied on measurements derived from entire tissue blocks, i.e. spanning all cortical layers and often including some underlying white matter (7). Laminar structure is a fundamental organizing principle of the cerebral cortex that relates to local micro-circuitry and inter-regional connectivity (20–23). In particular, inter-regional feedforward and feedback excitatory neural projections originate preferentially from supragranular (upper) layers II & III, and infragranular (lower) layers V & VI, respectively (23–27).

Spatial transcriptomics is a technique that can discriminate the expression levels of thousands of genes while maintaining positional information across tissue sections on a microscope slide (28, 29). With this technique it is possible to discern differences of transcriptomic profiles across cortical layers (30). It is known that laminar profiles of gene expression can vary between cortical regions (31), but although spatial transcriptomics has been applied to samples from the middle temporal cortex (30, 32), no such studies have yet included samples from the inferior frontal gyrus. Contrasting laminar profiles of gene expression between frontal and temporal regions of the core language network has therefore not been possible.

In the present study we applied 10x Genomics Visium spatial transcriptomics (28, 29) to discriminate laminar gene expression profiles in human cerebral cortical samples, derived from the left-hemisphere inferior frontal gyrus and superior temporal sulcus of three neurotypical adult donors (Figure 1). For human cerebral cortical tissue sections, each spot on a Visium spatial transcriptomic slide indexes the gene expression from an average of 3-5 cells (30). We used our data to identify genes that showed region-specific laminar expression profiles. Specifically, we sought differences between the frontal and temporal cortical regions in terms of supragranular layer II/III versus infragranular layer V/VI expression patterns. As connections between distant regions of the cerebral cortex are primarily made through excitatory projection neurons from these layers, we then integrated our spatial transcriptomic data with existing single-cell transcriptomic data from the human cerebral cortex, to identify genes that are expressed more highly in layer II/III and/or layer V/VI excitatory neurons than other cortical cell types, in addition to showing fronto-temporal differences in their infragranular versus supragranular expression patterns. We reasoned that the laminar expression patterns of such genes may be adapted to support frontal-temporal connectivity within the left-hemisphere language network.

**Figure 1.**
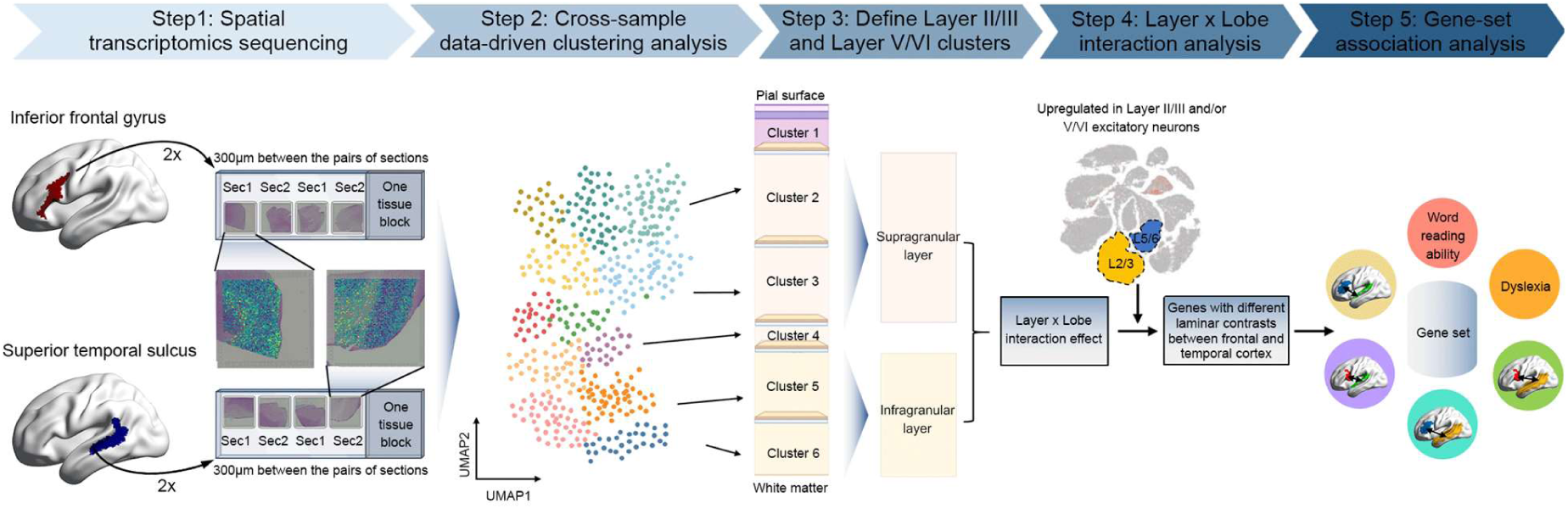
Schematic overview of the main steps in this study. <u>Step 1</u>: We acquired postmortem tissue sections from core regions of the left-hemisphere language network: the inferior frontal gyrus and posterior superior temporal sulcus, from three neurotypical donors. We performed spatial transcriptomics for each of 48 cortical tissue sections (3 donors x 2 regions x 2 blocks x 2 pairs of adjacent sections). <u>Step 2</u>: The resulting spot-level transcriptomic data were normalized to perform data-driven clustering analysis across all 48 sections, to identify layer-like clusters of spots in grey matter that matched across sections. <u>Step 3:</u> We annotated the data-driven clusters to cortical layers based on the spatial expression profiles of known layer marker genes, as well as classical cytoarchitecture. For each section, we pseudo-bulked the transcriptomic data by summing the unique molecular identifier counts across spots within layers II/III to define one supragranular bulked cluster, and within layers V/VI to define one infragranular bulked cluster. <u>Step 4</u>: We tested for genes showing layer*lobe interaction effects, i.e. genes whose supra- versus infragranular contrasts were different between the frontal and temporal cortex of the left-hemisphere. By reprocessing a previously published single-cell transcriptomic dataset generated from postmortem human cortical samples, we also identified genes that showed upregulation in layer II/III excitatory neurons and/or layer V/VI cortico-cortical projection neurons compared to other cortical cell types, as well as showing layer*lobe interactions. The laminar expression patterns of such genes may be adapted to support connectivity within the core language network. <u>Step 5</u>: The genes identified in step 4 were defined as a set, to test for a set-level association with white matter connectivity between core language-related brain regions, based on imaging genetics analysis in 30,814 individuals. We also tested for a set-level association with respect to word reading ability and dyslexia, based on previous large-scale genetic studies of these traits.

To test this hypothesis for the genes that we identified, we then analyzed genetic association data from a neuroimaging genetics study of 30,814 adults (33), with respect to structural connectivity of the main nerve fiber tract that links frontal and temporal regions of the core language network – the *arcuate fasciculus*. In addition, to test the possible relevance for language-related cognition of the genes we identified, we analyzed data from the largest genome-wide association studies of language-related traits performed to date: word reading ability in 33,959 individuals (34), and dyslexia in 51,800 adults who reported having a diagnosis versus 1,087,070 controls (35) (the latter based on data from 23andMe, Inc.).

## Results

### Data-driven cluster analysis based on gene expression defines cortical laminar structure

Separately for each of the three donors (Supplementary Table 1), tissue sectioning was performed using two blocks from the inferior frontal gyrus and two blocks from the posterior part of the superior temporal sulcus (Figure 1; Supplementary Figure 1). Separately for each block, we obtained two pairs of directly adjacent sections, resulting in a total of 48 sections (3 donors × 4 tissue blocks × 4 sections). After quality control (Methods), we measured expression levels for 22,170 genes across 140,192 spots on the Visium slides, corresponding to all 48 sections together (Supplementary Table 2, Supplementary Figures 2-3). Spots failing quality control in our data were especially concentrated in white matter, perhaps due to relatively low mRNA levels or diffusivity in the myelinated axons of neurons, while grey matter was well measured (Figure 2, Supplementary Figures 2-3).

**Figure 2.**
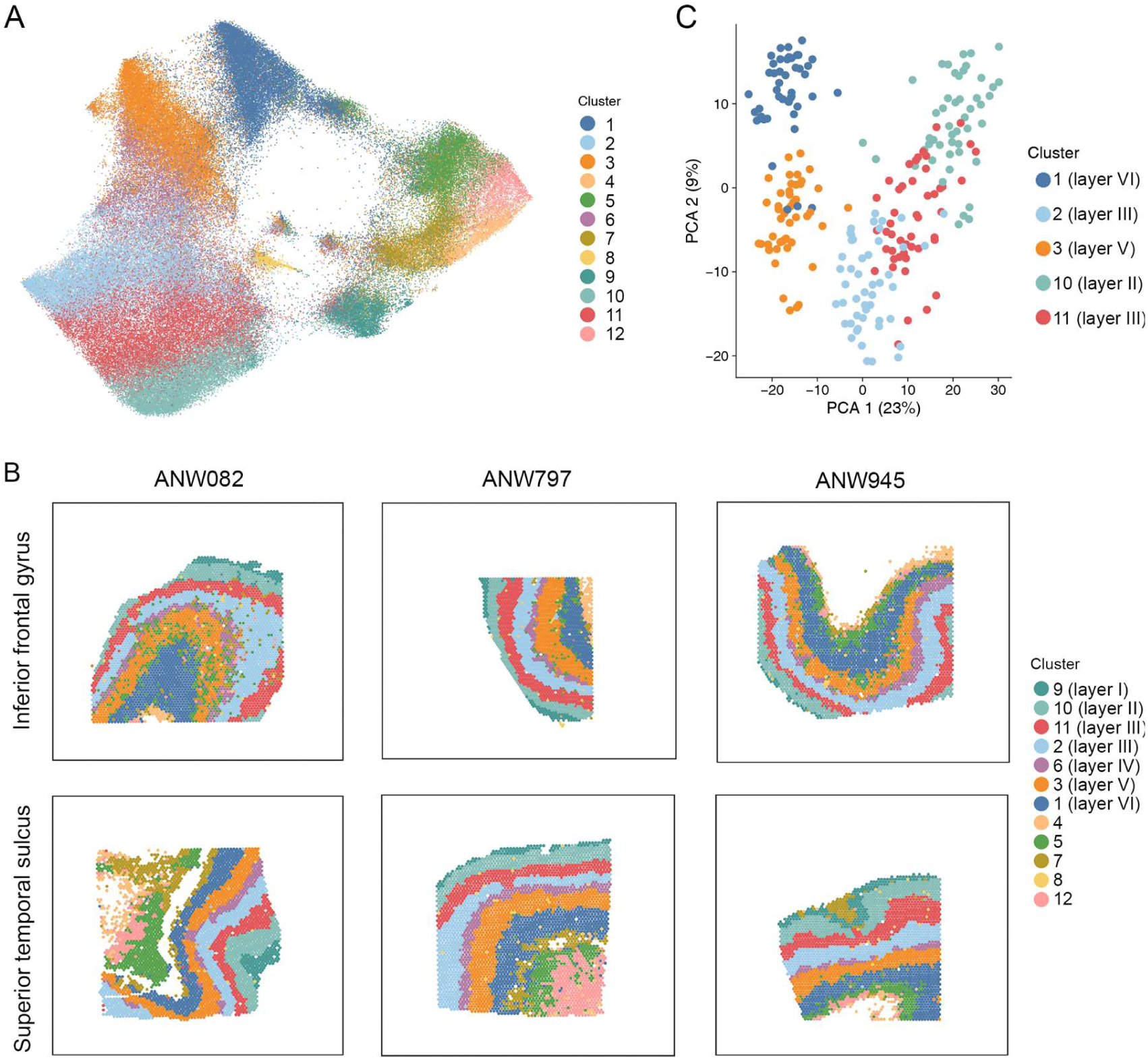
Data-driven analysis based on gene expression profiles identifies layer-like clustering in cerebral cortical tissue samples. (A) Uniform manifold approximation and projection mapping (UMAP) helps to visualize the similarities and differences between 12 data-driven clusters, based on combined analysis of data from 48 tissue sections from the inferior frontal gyrus and superior temporal sulcus. Each dot in the UMAP plot represents a single spatial transcriptomic spot from one of the 48 sections. Each spot captures transcriptomic expression from an average of 3-5 cells that correspond to that spot’s position on the slide. (B) Layer-like spatial patterns of data-driven clusters based on gene expression profiles, in six example sections of human cerebral cortex. For all 48 sections see Supplementary Figure 4. There were seven layer-like clusters in grey matter, and five clusters that were located in white matter or sporadically located without laminar appearances. Cluster-layer correspondence was determined through known layer marker genes and cytoarchitecture, whereby layer I = cluster 9; layer II = cluster 10; layer III = clusters 2 &11; layer IV = cluster 6; layer V = cluster 3; layer VI = cluster 1 (see main text). (C) Principal component analysis (PCA) of pseudo-bulked data from five clusters corresponding to cortical layers II, III, V and VI. The first principal component (PCA 1) explained 23% of variance in the transcriptomic data (as indexed from the top 10% of variable genes) and supported a primary distinction of supragranular from infragranular clusters in terms of transcriptomic profiles. The second principal component (PCA 2) explained 9% of the variance and reflected proximity to the granular layer.

Data were normalized and harmonized for sample effects (Methods). We then performed clustering analysis using the data from all sections together, using BayesSpace (36). This approach identified corresponding data-driven clusters of spots across sections that were matched based on their gene expression profiles. The main data-driven clusters had clear layer-like appearances in all sections (Figure 2B, Supplementary Figure 4). UMAP analysis confirmed that the top components of variation in the transcriptomic data were related to layer-like differences and the grey-white matter distinction (Fig. 2).

We then annotated the data-driven clusters with reference to known layer marker genes (30, 37, 38) and structural cytoarchitecture visible by Nissl staining (Methods, Figure 3A, Supplementary Figures 5-6). The cerebral cortex is classically divided into six layers (39, 40). *AQP4*, *FABP7* and *RELN*, as markers of layer I, showed highest expression in cluster 9 that was defined at the outermost surface (Figure 2B, Supplementary Figure 5). Directly underlying this, cluster 10 was enriched for the expression of layer II markers, such as *ENC1* (Supplementary Figure 5). Our data-driven analysis based on transcriptomic profiles distinguished two clusters corresponding to layer III (clusters 2 & 11), and both of these clusters showed higher expression of layer III marker *ADCYAP1* than other clusters (Supplementary Figure 5). *RORB* as a canonical marker of layer IV, was expressed most highly in cluster 6 (Supplementary Figure 5). Layer V marker *TRABD2A* was expressed most highly in cluster 3 (Supplementary Figure 5), and layer VI marker *CCK* was expressed most highly in cluster 1 (Supplementary Figure 5). All of these marker-based annotations matched the expected order of spatial layering from upper to lower layers. Additional clusters were located in white matter, with higher expression of marker *MBP* than other clusters (Supplementary Figure 5), or were based on a smaller number of sporadically distributed spots (Figure 2B, Supplementary Figure 4).

**Figure 3.**
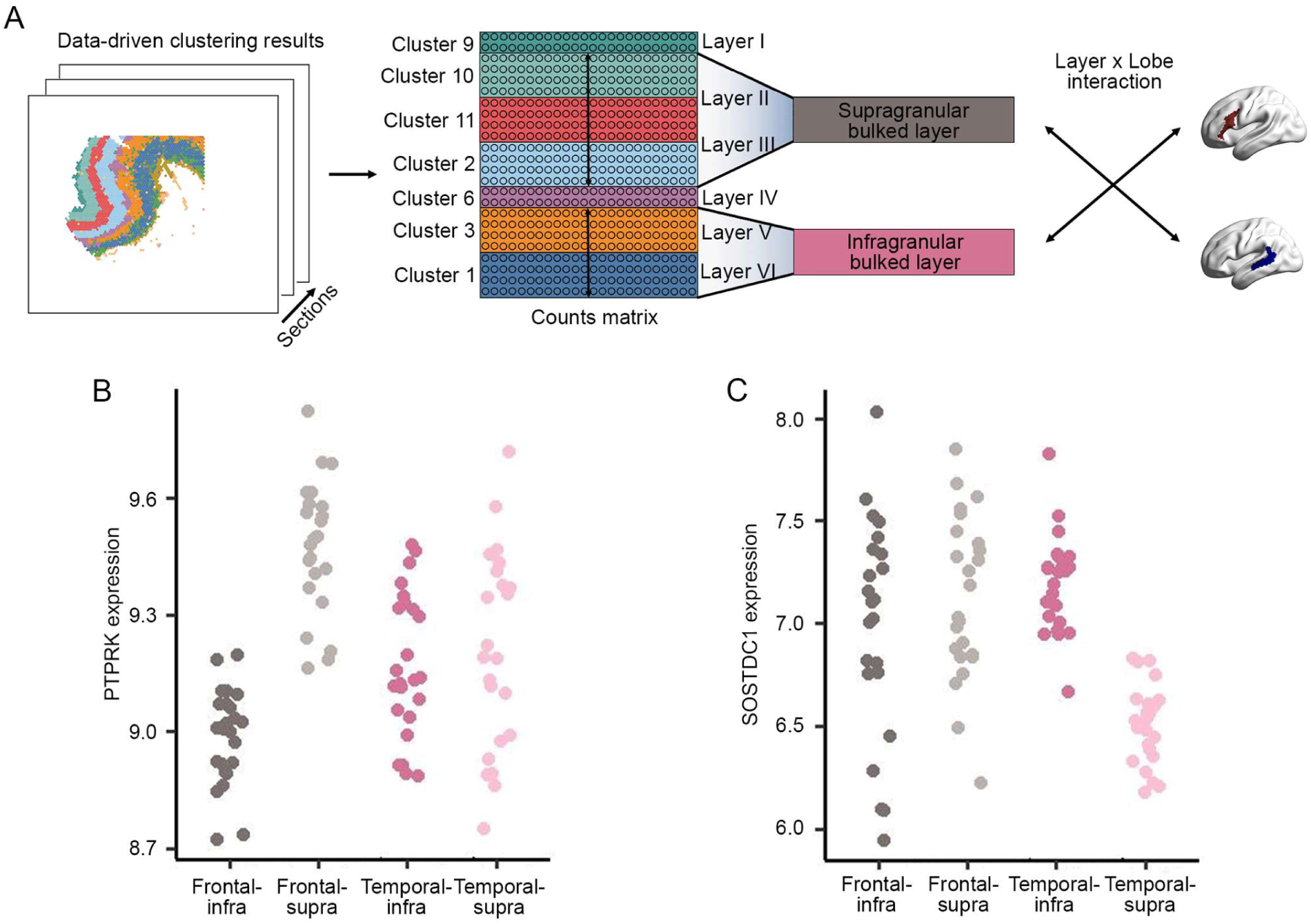
Identiying genes with supra- versus infragranular contrasts that differ between frontal and temporal regions of the left-hemisphere language network. (A) Schematic of the pseudo-bulking procedure that integrated the spatial transcriptomic data from spot-level to layer-level within each cortical tissue section, separately for supragranular layers II/III versus infragranular layers V/VI. The pseudo-bulked data were then used in linear mixed effects modelling to test each gene for a layer*lobe interaction effect (supra- vs infragranular)*(frontal vs temporal), while controlling for donor, tissue block and adjacent sections. (B & C) Expression data for two genes with the most significant layer*lobe interaction effects, where each dot represents one of 48 sections for each lobe and layer combination: (B) *PTPRK* showed markedly higher supragranular than infragranular expression in the inferior frontal gyrus, but supragranular and infragranular levels were similar in the superior temporal sulcus. (C) *SOSTDC1* showed similar supragranular and infragranular levels in the inferior frontal gyrus, but markedly higher infragranular than supragranular expression in the superior temporal sulcus. For comparable plots for all 56 genes that showed significant layer*lobe interaction effects as well as upregulation in layer II/III and/or layer V/VI excitatory neurons compared to other cortical cell types, see Supplementary Figure 7.

### Supragranular versus infragranular layers

As excitatory neural projections from supragranular layers II & III, and infragranular layers V & VI, are primarily responsible for inter-areal projections (23), we focused on these four layers for our subsequent analyses. We therefore left aside data from layer I (i.e. cluster 9), and the granular layer IV (named for its granular appearance under light microscopy) that separates the supragranular from infragranular layers. This meant focusing on data from five clusters: supragranular cluster 10 (layer II), supragranular clusters 2 & 11 (layer III), infragranular cluster 3 (layer V) and infragranular cluster 1 (layer VI) (Figures 2B, 3A).

Separately for each of the 48 tissue sections and each of these five clusters, we performed “pseudo-bulking” by summing and normalizing the unique molecular identifier counts across all spots for each gene (Methods), to generate a single section- and cluster-specific expression value per gene (therefore 240 values per gene, from 48 sections x 5 clusters). Principal component analysis based on the 10% of most variable genes (Methods) showed how the sections and clusters related in terms of transcriptional profile similarity (Figure 2C). The first component explained 23% of the variance and distinguished supragranular clusters 1 & 3 from infragranular clusters 2, 10 & 11 (Figure 2B, 2C). The second component explained 9% of the variance and captured laminar differentiation in a manner that reflected proximity to the granular layer (Figure 2B, 2C). These findings further confirmed that cortical layers can be meaningfully defined based on transcriptional profiles, and supported a primary distinction of supragranular layers II/III from infragranular layers V/VI at the mRNA level. The findings also indicated that the stereotypical layer arrangement, as reflected by overall transcriptional profiles, is conserved between the inferior frontal gyrus and superior temporal sulcus of the left-hemisphere. We therefore expected that any inter-areal differences of laminar gene expression between these regions would be relatively subtle, and/or restricted to a minority of genes.

### Identifying genes that show different laminar expression patterns between frontal and temporal language cortex

For the following analyses we retained 12,656 genes for which canonical transcripts were defined according to the human reference genome GRCh38. For each of the 48 tissue sections separately, we pseudo-bulked data from spots in clusters 2, 10 & 11 to create a single supragranular layer II/III bulked cluster, and from clusters 1 & 3 to create a single infragranular layer V/VI bulked cluster (Methods). This resulted in 96 expression values per gene (48 sections x 2 bulked clusters). We then used linear mixed effect modeling to test for genes that showed layer*lobe interaction effects, i.e. genes whose supra- versus infragranular contrasts were different between the inferior frontal gyrus and superior temporal sulcus (Figure 3). We controlled for donor and tissue block as fixed effects, while also accounting for adjacent sections in the model (Methods).

Seventy-two genes showed significant layer*lobe interaction effects at FDR p<0.01 (Supplementary Table 3). Two genes particularly stood out in terms of statistical significance: *PTPRK* (t=9.20, p=1.05×10^-14^) and *SOSTDC1* (t=8.58, p=2.07×10^-13^) (Supplementary Table 3) (Figure 3B, 3C). In the inferior frontal gyrus, *PTPRK* showed markedly higher supragranular than infragranular expression, whereas in the superior temporal sulcus this gene showed similar expression in supragranular and infragranular layers, thus giving rise to a layer*lobe statistical interaction (Figure 3B). *PTPRK* encodes a signaling molecule of the protein tyrosine phosphatase family. For *SOSTDC1*, there was markedly higher infragranular than supragranular expression in the superior temporal sulcus, while in the inferior frontal gyrus the expression level was similar in infragranular and supragranular layers (Figure 3C). *SOSTDC1* encodes a secreted protein of the sclerostin family that functions as an antagonist to growth factors of the bone morphogenetic protein (BMP) family, and also affects Wnt signalling (41). The 72-gene set showed enrichments at FDR p<0.01 for thirteen biological processes defined in the Gene Ontology (42), most significantly for functions related to neuron projection/axon guidance, driven by the genes encoding signalling molecules NELL1, PTPRM, SLIT1 and SLIT2 (Supplementary Table 4) (43). Other implicated processes included ion transport and regulation of catecholamine secretion (Supplementary Table 4).

### Genes upregulated in layer II/III or layer V/VI excitatory neurons

To focus on the laminar profiles of genes expressed in the cells that primarily form inter-areal projections, we then used existing single-cell RNA sequencing data to identify genes with higher expression in excitatory neurons of layers II/II and/or layers V/VI compared to other cortical cell types. We reprocessed single cell data derived from the dorsolateral prefrontal cortex and anterior cingulate cortex from 31 donors (44) (Methods). This dataset included post-quality control gene expression data from 104,559 nuclei that had been annotated to 17 defined cell types. For each of 12,155 genes that overlapped with those in our spatial transcriptomics dataset, we extracted and processed single cell gene expression data to aggregate unique molecular identifier counts for each of the 17 defined cell types (44) (Methods). These profiles were then used for differential expression analysis to identify genes that were expressed more highly in the cell type ‘layer II/III excitatory neurons’ and/or the cell type ‘layer V/VI cortico-cortical projection neurons’ compared to all other cell types combined (see Methods). At FDR p<0.01 there were 2219 genes upregulated in layer II/III excitatory neurons, and 1732 genes upregulated in layer V/VI cortico-cortical projection neurons (Supplementary Tables 5 and 6), which yielded a union list of 2622 genes showing upregulation in one or both of these cell types.

Among the 2622 genes, there were 56 genes that showed significant layer*lobe interactions at FDR p<0.01, i.e. 56 genes with supra- versus infragranular contrasts that differed between the inferior frontal gyrus and superior temporal sulcus, in addition to showing upregulation in excitatory neurons of layer II/III and/or layer V/VI cortico-cortical projection neurons (Supplementary Figure 7; Supplementary Table 7). These again included *PTPRK* and *SOSTDC1* as the two most significant genes with layer*lobe interaction effects (Supplementary Table 7), and gene ontology enrichment analysis again pointed most significantly to neuron projection/axon guidance, driven by *NELL1, NELL2, SLIT1* and *SLIT2* (Supplementary Table 8). Magnesium ion responsiveness was also implicated, driven by *RYR3* and *SNCA* (Supplementary Table 8). Other individual genes among the set of 56 included the transcription factors *LMO3* and *LMO4;* the latter might be involved in the patterning of fetal brain left-right asymmetry (45), and *CDH10* encoding a cell adhesion molecule of the cadherin family that is involved in layer and circuit formation (46) (Supplementary Table 8). All 56 genes that showed upregulation in excitatory neurons of layer II/III and/or layer V/VI cortico-cortical projection neurons, together with significant layer*lobe interactions at FDR p<0.01, can be found in Supplementary Table 7.

### Genetic association analysis with respect to brain and behavioural variability

A previous genome-wide association study in 30,810 adults from the UK Biobank dataset mapped associations of common genetic variants in the population with individual differences in brain-wide structural connectivity, as determined by tractography based on diffusion imaging (33). We made use of the genome-wide summary statistics from that study, which index the extent to which each common variant in the genome is associated with a given structural connection in the brain. For the present study, we extracted data for four frontal-temporal connections that link the approximate left-hemisphere regions from which we sampled postmortem brain tissue for spatial transcriptomics: i.e. between the *pars opercularis* and superior temporal cortex, *pars triangularis* and superior temporal cortex, *pars opercularis* and middle temporal cortex, and *pars triangularis* and middle temporal cortex (Figure 1, Supplementary Figure 1, Supplementary Figure 8). These white matter connections are primarily via the *arcuate fasciculus* (33) (Supplementary Figure 8).

We focussed on the 56 genes that showed significant evidence in the present study for having different laminar patterns between frontal and temporal cortex, and upregulation in excitatory neurons of layer II/III and/or layer V/VI cortico-cortical projection neurons compared to other cortical cell types, i.e. the neurons that primarily form inter-areal connections. We applied the Gene-set Association analysis Using Sparse Signals (GAUSS) software (47), which tests for gene-set level association based on genome-wide summary statistics, and also identifies the specific genes that drive a set-level association. The 56 genes showed a significant set-level associaton with white matter connectivity between the *pars triangularis* and middle temporal cortex (unadjusted p=0.004, Bonferroni-adjusted p=0.016 for testing four connectivities) (Supplementary Table 9). GAUSS identified 26 genes driving this significant association, with 6 genes individually at p<0.05: *BHLHE22*, *COL5A2*, *NELL2*, *RYR3, SLIT1*, *SLIT2* (Supplementary Table 9). Connectivity between the *pars triangularis* and superior temporal cortex also showed nominally significant set-level association with the 56 genes (unadjusted p=0.02), but this did not survive Bonferroni correction.

In terms of language-related behavioural traits, by far the largest genome-wide association studies performed to date have been of word reading ability in 33,959 individuals (34) and dyslexia in 51,800 adults who reported having a diagnosis versus 1,087,070 controls (35). We tested the 56 genes for a set-level association with word reading ability and separately also with dyslexia, based on the genome-wide association summary statistics from those studies, for SNPs spanning the genomic locus of each gene plus 50 kilobases upstream and downstream (34, 35). The 56 genes showed no significant set-level association with word reading ability, but a highly significant association with dyslexia (p=6.22×10^-9^) (Supplementary Table 10). GAUSS identified 24 genes driving the association with dyslexia, with 14 genes individually at p<0.05: *BHLHE22*, *CDH10*, *DAB1*, *DIAPH1*, *FBXO32*, *GABRD*, *GPR26*, *KCNH5*, *KIRREL3*, *NEFH*, *OXR1*, *SLIT1*, *SLIT2, SNCA* (Supplementary Table 10). Three of these 14 genes were also among the 6 associated with structural connectivity between the *pars triangularis* and middle temporal cortex, suggesting that region-specific laminar expression patterns of these genes might be especially important for supporting language-related cognition: *BHLHE22* which encodes a neural transcription factor, and *SLIT1* and *SLIT2* which encode axon guidance molecules.

## Discussion

In this study we created a new spatial transcriptomics dataset from postmortem human cerebral cortical tissue, focused on core regions of the left-hemisphere language network. We identified genes with region-specific laminar expression profiles, i.e. that showed supragranular versus infragranular contrasts that differed between the inferior frontal gyrus and superior temporal sulcus. In combination with single cell transcriptomic data, there were 56 genes that additionally showed upregulation in layer II/III excitatory neurons and/or layer V/VI excitatory cortico-cortical projection neurons - the cell types that are primarily responsible for inter-areal connections - compared to other cortical cell types. Consistent with this, the 56 genes showed a set-level association with frontal-temporal white matter connectivity, and also with the language- and reading-related disorder dyslexia, in large-scale genome-wide association data. Genes involved in axon guidance were especially implicated by these analyses, including *SLIT1* and *SLIT2*.

Spatial transcriptomic data-driven clustering revealed robustly matching layers across cortical tissue sections from frontal and temporal regions, which indicates that laminar patterns are in fact largely similar between the inferior frontal gyrus and superior temporal sulcus, in terms of overall gene expression profiles. However, we aimed to identify genes that do not conform to this overall homogeneity in the present study. This was because language-related cognition is likely to be supported by specialized areal functions, and uniquely adapted inter-areal connectivity, between frontal and temporal regions of the left hemisphere (in addition to pan-cortical aspects of laminar organization). Our findings confirm the existence of regional variation in laminar expression profiles for a minority of genes, following pioneering work based on a smaller number of genes and other cortical regions (31). These findings therefore establish regional variation of laminar gene expression as an organizing principal in the human cerebral cortex.

Invasive, *in vivo* axon tracing, especially in the visual system of macaque monkeys, has shown that inter-regional feedforward and feedback connections tend to originate from upper and lower cortical layers respectively (23, 24). Clearly such invasive tracing methods cannot be applied in humans, which means that layer-specific frontal-temporal connections within the human language network have not been directly observed. Nonetheless, the frontal and temporal regions of the core language network are connected by prominent white matter tracts, notably the *arcuate fasciculus* that has undergone specific left-lateralized anatomical changes in humans compared to macaques (48). This is consistent with regional and network-level specialization of the frontal and temporal regions in the left-hemisphere, to support language-related cognition. The present study provides indirect evidence that inter-areal connections within the left-hemisphere language network are supported by region-dependent laminar profiles, because genomic variants within the implicated genes showed association with inter-individual variation in frontal-temporal white matter connectivity.

However, we cannot exclude that genes with regional variation of laminar expression may affect inter-areal connectivity through other mechanisms, not only related to their region-dependent laminar expression profiles. It also remains unclear to what extent feedforward and feedback connectivity may operate between the frontal and temporal regions of the left-hemisphere language network. These mechanisms are most easily understood with respect to bottom-up and top-down transmission of information between regions on different hierarchical levels, from sensory cortex to association cortex. The inferior frontal gyrus and superior temporal sulcus are both implicated in high-level linguistic functions, including sentence-level processing (1–3). Therefore these regions may be broadly comparable in terms of hierarchy within the language network. Layer-specific functional MRI has been applied to dissociate top-down and bottom-up signal contributions to the left occipitotemporal sulcus during word reading (49), but this region does not overlap with the temporal region analyzed in the present study.

The genes that we implicated in this study showed their most significant Gene Ontology set-level enrichment for functioning in axon guidance. For example, SLIT1 and SLIT2 are well known as extracellular axon guidance molecules that are involved in the patterning of brain networks during development (43), as well as other aspects of neocortical formation (50). Less is known of the roles of these genes in mature adult brains, long after inter-areal nerve fiber connections are established. They may have roles in the regulation of tissue integrity and homeostasis (51, 52), through mechanisms similar to their developmental roles, for example in regulating cell-cell adhesion. Consistent with this, SLIT proteins have been implicated in synaptogenesis through forming complexes with presynaptic Neurexin and postsynaptic Robo proteins (53), and their expression in the adult cortex may therefore affect synaptic plasticity and stability. Our data suggest that such roles contribute to cortical regional and laminar specialization of function.

As regards dyslexia, this disorder of reading is closely linked to linguistic cognition in many affected individuals, including reduced phonological awareness and sometimes language impairment (54–56). Dyslexia has also been linked to altered frontal-temporal structural connectivity via the *arcuate fasciculus*, although results on this have been inconsistent across studies (57, 58). While reading is a cultural innovation, it recruits much of the neural circuitry that underpins the human capacity for oral language (59). The present study found that genes with laminar expression patterns that differ between the frontal and temporal language cortex, and are also upregulated in excitatory neurons of layers II/III and/or layers V/VI, show a highly significant set-level association with dyslexia. For this analysis we used data from a genetic study of 51,800 adults who reported having had a diagnosis, versus 1,087,070 controls (35). It is likely that many of the 51,800 individuals received their diagnoses during childhood, although information on this was not recorded, and some may have received diagnoses in adulthood. For *SLIT* genes and other genes involved in axon guidance, their contributions to dyslexia may occur to a large extent during neurodevelopment, but perhaps also via functions in the adult brain as discussed above, consistent with the neural expression of such genes in the adult cerebral cortex.

The lack of a gene set-level association with word reading ability may have been due to the smaller sample size of the genome-wide association study from which we used summary statistics for that trait (33,959 individuals with quantitative data on word reading skills) (34), compared to the larger study of dyslexia (∼1.14 million individuals) (35). Future studies would benefit from larger-scale GWAS of additional language-related traits such as expressive and receptive language abilities.

Transcription from genomic DNA to mRNA occurs in the cell nucleus, and translation from mRNA to proteins occurs prominently in the cell body, from where proteins can be transported along axons and dendrites to other locations where they are needed - for example presynaptic boutons. However, movement of mRNA molecules along axons and dendrites also occurs, to support local translation distantly from the cell body (60). In the present study we interpreted laminar gene expression profiles as reflecting primarily cell body gene expression, which for inter-areal projection neurons implies the origins of connections, rather than their terminals. Also we used single-nucleus gene expression data to inform our analyses, which necessarily misses mRNA in axons and dendrites. In adult brain tissue, the extent to which neural translation occurs locally in the neuropil versus in the cell body is uncertain (60), but translation in the neuropil may be a substantial contributor for some proteins (61). With the spatial transcriptomic technology used in the present study, each spot has previously been shown to measure mRNA molecules from an average of 3-5 cell bodies in cerebral cortical tissue sections, but a contribution from neuropil is also expected, and a minority of spots that are not overlain by cell bodies may measure neuropil mRNAs only (30). In this case, laminar differences of gene expression may relate partly to the terminals of inter-areal connections, as well as the abundances of different cell types or the levels of expression within those cell types.

To summarize, this study made several contributions to understanding the cerebral cortical infrastructure underlying language. We identified genes with region-dependent laminar expression profiles that may support regional functional specialization. Ours is among a small number of studies to have shown cortical regional variation in laminar gene expression, and helps to establish this as an organizing principal of the human brain more generally. Through additionally focusing on upregulation of transcription in excitatory projection neurons, our study identified genes that may affect inter-areal connectivity within the left-hemisphere language network. This possibility was supported by genetic association data that implicated several of the relevant genes in fronto-temporal white matter connectivity, and the language-related trait dyslexia.

## Methods

### Donors and sampling

This study was approved by the Ethics Committee Faculty of Social Sciences, Radboud University Nijmegen, and the ethics and Biobank Review Committee, Free University Medical Center Amsterdam. All donors gave written informed consent to the human body bequest program at the department of Anatomy and Neurosciences, Amsterdam UMC – location VUmc (governed by the human tissue act (‘ter beschikking stelling’, Artikel 18, lid 1 en 19 van de Wet op de Lijkbezorging, 1991)). Collection of brain tissue was facilitated though the Normal Aging Brain Collection Amsterdam (NABCA) biobank. The standardized protocol at NABCA includes craniotomy and dissection of left hemisphere tissue blocks followed by freezing (liquid nitric oxygen), with a postmortem delay to autopsy of around 8 hours (8). For the present study we obtained postmortem brain tissue from three neurotypical donors: a 59-year-old male, a 59-year-old female, and a 63-year-old female (Supplementary Table 1). All three postmortem brains were confirmed to have minimal pathology by macro- and microscopic investigation, and RNA integrity numbers of at least 7 (Supplementary Table 1) (8, 62). From each donor we sampled from four left-hemisphere tissue blocks of roughly 1.5cm^3^ each (containing both grey and white matter): two blocks from the inferior frontal gyrus and two blocks from the posterior superior temporal sulcus (see Supplementary Figure 1 for the sampled regions).

### Sectioning and spatial transcriptomic data generation

From each tissue block we took two pairs of adjacent 10μm-thick tissue sections that would be used for spatial transcriptomics, resulting in a total of 48 tissue sections: 3 donors x 2 lobes (frontal and temporal) x 2 blocks x 2 adjacent pairs of sections. The section pairs were separated by 300μm (Figure 1). The sectioning was performed within a cryostat (Thermo Fisher NX70) following the manufacturer’s protocol (Visium Tissue Preparation Guide, 10x Genomics CG000240 RevA), and aimed to span across the cortical layers. Frozen 10μm-thick tissue sections were mounted on Tissue Optimization Slides (Visium Spatial Tissue Optimization Slide & Reagent Kit, 10x Genomics 1000193) and Gene Expression Slides (Visium Spatial Gene Expression Slide & Reagent Kit, 10x Genomics, 1000184) and stored at -80°C until further processing. Additional sections from within the 300μm interval were taken for other parts of the study (see below) and stored at -80°C until use.

Permeabilization time was optimized following the manufacturer’s protocol (Visium Spatial Tissue Optimization, 10x Genomics CG000238 RevA). Fluorescence images of sections for tissue optimization were taken using a AxioScan Z1 SlideScanner (Zeiss) with Hamamatsu Orca Flash camera, a Cy5 filter, a Plan-Apochromat 20x/0.8 M27 objective and Zeiss Zen v.2.6 (Blue Edition) software. A permeabilization time of 18-minutes was used for the Visium Spatial Gene Expression workflow. Frozen tissue sections on gene expression slides were processed for spatial transcriptomics using the Visium Spatial Gene Expression Slide & Reagent Kit (10x Genomics, 1000184) following the manufacturer’s protocols (Visium Spatial Gene Expression User Guide,10x Genomics CG000239 RevA). Briefly, brightfield images of H&E-stained sections were acquired using an AxioScan Z1 slide scanner (Zeiss) with a Hitachi HV-F292SCL camera, a Plan-Apochromat 20x/0.8 M27 objective and Zeiss Zen v.2.6 (Blue Edition) software. H&E-stained sections were then permeabilized for 18 minutes, followed by reverse transcription, second-strand synthesis, denaturation, cDNA amplification and quality control, and library construction. qPCR was performed using the KAPA SYBR FAST kit (KAPA Biosystems) and BioRad CFX96 Real-Time System. The cDNA amplification cycle number was determined at ∼25% of the peak fluorescence value. The final libraries were processed and a BioRad T100 Thermal Cycler was used for PCR reactions during library construction. The final libraries were sequenced on a NovaSeq 6000 System (Illumina) for 150 bp pair-end reads. The sequencing reads were then trimmed to read 1: 28 cycles; i7 index read: 10 cycles; i5 index read: 10 cycles; and read 2: 91 cycles, followed by demultiplexing using Space Ranger software v.1.2.2 (10x Genomics). The 48 libraries were sequenced to a median depth of 469×10^-6^ reads.

### Spatial transcriptomic data processing and quality control

We processed the raw FASTQ files and H&E histology images of sections with the Space Ranger software v.1.2.2, using STAR v.2.5.1b (63) for alignment against the Cell Ranger reference genome refdata-cellranger-GRCh38-3.0.0, available at: http://cf.10xgenomics.com/supp/cell-exp/refdata-cellranger-GRCh38-3.0.0.tar.gz. H&E histology images were rotated and resized to 2000 x 2000 pixels using FIJI/ImageJ (v.1.53t). Default parameters for spaceranger count v.1.2.2 was used to generate count matrix files and QC metrics (https://support.10xgenomics.com/spatial-gene-expression/software/pipelines/latest/using/count). QC metrics returned by this software are available in Supplementary Table 2.

This processing resulted in unique molecular identifier (UMI)/feature-barcode matrices for each of the 48 sections. We read the raw feature-barcode matrix from each section, coupled with its corresponding histology image, to construct a customized object using the SummarizedExperiment R package (64). The data across all 48 sections were then aggregated to form one SingleCellExperiment object (65). There were 170,157 detected spots corresponding to all 48 sections together, with a mean of 6783 UMIs per spot, and a mean of 2782 genes per spot. We excluded spots according to spot-wise quality control metrics using the default settings of the perCellQCMetrics and quickPerCellQC functions from the Scran v1.18.7 R Bioconductor package, which considers the log-total UMI count, log-number of detected features, and percentage of counts in specified ‘control’ gene sets (mitochondrial genes, spike-in transcripts) (66). We also excluded spots in parts where the sections had folded over on the spatial transcriptomic slides (Supplementary Figure 3). We then excluded genes that were expressed in less than 0.01% of spots across all of the 48 samples combined, as well as 13 mitochondrial genes. These steps left 140,192 spots from which a total of 22,170 genes were measured.

### Data-driven clustering to match across sections

We used the functions quickCluster and computeSumFactors within Scran (66), and the logNormCounts function within the Scater package v1.18.6 (67), to compute log normalized gene expression counts at the spot level. The normalized data were then used to fit a model with respect to gene mean expression and variance using the modelGeneVar function of Scran (66), followed by identifying the top 10% of variable genes. This set of highly variable genes was used to compute principal components using the runPCA function within Scater (67), generating the top 50 components. We used Harmony (68), an algorithm that can perform integration of gene expression data from multiple spatial transcriptomics datasets, to adjust for batch effects on these 50 components.

Harmonized components were then used to perform clustering analysis with BayesSpace (36), a deep-learning and Bayesian algorithm that identifies clusters of spots based on transcriptomic profile similarity, but also gives higher weight for clustering to physically close spots. We performed clustering across all 48 sections in a single analysis, which had the advantage of identifying matching layer-like clusters across sections, regardless of variability in terms of the exact orientation of sectioning across layers (Figure 2B, Supplementary Figure 4). We set the clustering implementation to return twelve clusters, which identified seven layer-like clusters within grey matter that were consistently located across sections, and five clusters that were located either in white matter or sporadically distributed without layer-like appearances, the latter based on small numbers of spots (Figure 2, Supplementary Figure 4). In this way we were confident to have identified the main layer-like clusters within grey matter, which were the focus of our study. The outermost layer-like cluster, i.e. cluster 9 (corresponding to cortical layer I on the basis of marker gene expression and histological analysis – see below), was not identified in all 48 sections (Supplementary Figure 4), probably due to surface tissue loss during freezing/handling/sectioning.

We also generated a two-dimensional map based on uniform manifold approximation and projection (UMAP) with the Scater package (69) across all spots, to visualize how the clusters compared to each other in terms of transcriptional similarity (Figure 2A).

### Mapping data-driven clusters to cortical layers

We extracted the expression data for 12 genes that have been reported as robust layer markers in human cerebral cortical tissue (30), and plotted them separately across each of the twelve clusters (Supplementary Figure 5). This revealed the expected pattern of upper-to-lower layer identities as described in the Results section (above), whereby the following correspondences were identified: layer I = cluster 9; layer II = cluster 10; layer III = clusters 2 &11; layer IV = cluster 6; layer V = cluster 3; layer VI = cluster 1 (Figure 2B, Figure 3A, Supplementary Figure 5).

### Cresyl-violet/Nissl staining

In addition, we used extra sections taken from the 300μm intervals between spatially adjacent section pairs from three tissue blocks, to perform Cresyl-violet/Nissl staining. Fresh-frozen sections (10 µm) adjacent to the section used for Visium spatial transcriptomics was used for Cresyl-violet/Nissl staining. Sections were first fixated in 4% paraformaldehyde (in PBS, pH=7.4) for 10 minutes at room temperature. Fixated sections were washed twice for 5 minutes each with 1x PBS, followed by serial hydration in 70% ethanol, 50% ethanol and demi water, 2 minutes each. Sections then went through rapid dehydration in 70%, 96% and 100% ethanol, and were air-dried. Air-dried sections were stained in 0.1% Cresylviolet (Sigma) in MilliQ water at 56°C for 20 minutes, then incubated in 96% ethanol for 1 minute to remove excess staining. Stained sections were dehydrated in 96% ethanol for 2 minutes twice, in 100% ethanol for 2 minutes three times, in xylol (Sigma) for 2 minutes three times, and mounted in Entellen New (Sigma). Brightfield images were acquired using a AxioScan Z1 SlideScanner (Zeiss) with a Hitachi HV-F292SCL camera, a Plan-Apochromat 20x/0.8 M27 objective and Zeiss Zen v.2.6 (Blue Edition) software. Images were analysed with FIJI/ImageJ (v.1.53t) and QuPath0.2.3. Trained experimenters from different institutes (authors SvH, LEJ) then independently confirmed layer identification according to the classical six-layer schema, based on structural cytoarchitecture and manual boundary definitions (Supplementary Figure 6).

### Supragranular versus infragranular expression

We focused our subsequent analyses on supragranular layers II/III (clusters 2, 10, 11) and infragranular layers V/VI (clusters 1, 3). For each gene, we pseudo-bulked the spot-level data into cluster-level data (70–72), by summing the raw gene-expression counts across all spots in a given section and cluster, resulting in a new SingleCellExperiment object containing 22,170 genes across 240 clusters (i.e. 48 sections × 5 clusters). This pseudo-bulking procedure reduced sparsity and increased the coverage of genes compared to spot-wise data. Using the same software and functions as described above with respect to the processing for data-driven clustering analysis, we normalized the pseudo-bulked, cluster-enriched gene expression matrix, identified the top 10% of variable genes, and computed the top 2 principal components to visualize how the 48 sections x 5 clusters relate in terms of transcriptome profile similarity. This supported a primary distinction between the supragranular and infragranular clusters (Figure 2C), and we therefore repeated the pseudo-bulking, but now for spots assigned to supragranular clusters 2, 10 and 11 all together, and spots assigned to infragranular clusters 1 and 3 all together, resulting in a new SingleCellExperiment object that comprised gene expression data from 48 sections x 2 pseudo-bulked clusters. We only retained genes with canonical transcripts defined in reference human genome GRCh38, https://ftp.ncbi.nlm.nih.gov/refseq/MANE/MANE_human/release_0.93/, for a total of 12,656 genes.

With the normalized pseudo-bulk profiles, we computed the correlation structure between spatially adjacent sections using the duplicateCorrelation function of the Limma software v3.46.0 (73). We then applied linear mixed effect modelling using the lmFit and eBayes functions of Limma (73), to test a model for each gene whereby its expression level varied depending on the main effect of layer (supra- versus infragranular expression, i.e. a binary variable), lobe (frontal versus temporal, another binary variable), the layer*lobe interaction (our primary interest was to find genes with different laminar patterns between the frontal and temporal regions), with donor and tissue block as fixed effects, and blocking the 24 pairs of spatially adjacent sections according to their correlation coefficient matrix. In fitting such a model, Limma uses data from all genes to improve the variance estimate for any given gene, which helps to overcome relatively small sample sizes in transcriptomic studies (73). FDR correction was applied at adjusted P<0.01 to identify genes with significant layer*lobe interaction effects. The list of genes with FDR-adjusted P<0.01 was input to the software Enrichr (74) (as implemented at https://maayanlab.cloud/Enrichr/) for a descriptive Gene Ontology analysis.

### Cell-type analysis

We next made use of an existing single-cell transcriptomic dataset based on 104,559 nuclei from 41 postmortem tissue samples from the dorsolateral prefrontal cortex or anterior cingulate cortex of 15 individuals with autism and 16 neurotypical controls (44). That study had used the single-cell gene expression data to define 17 major cell types: fibrous astrocytes, protoplasmic astrocytes, endothelial, parvalbumin interneurons, somatostatin interneurons, SV2C interneurons, VIP interneurons, layer II/III excitatory neurons, layer IV excitatory neurons, layer V/VI corticofugal projection neurons, layer V/VI cortico-cortical projection neurons, microglia, maturing neurons, NRGN-expressing neurons I, NRGN-expressing neurons II, oligodendrocytes and oligodendrocyte progenitor cells. We aggregated UMI counts from all 104,559 nuclei to create cell type-specific log-transformed normalized counts for each of the 17 cell types, resulting in a set of pseudo-bulked profiles for all unique donor-region-cell type combinations. For each gene, we then used its pseudo-bulked profiles to test for upregulation in a given cell type compared to all other cell types, using the lmFit and eBayes functions in Limma (73), while adjusting for the fixed effects of brain region, age, sex and diagnosis, and a random intercept of donor. We then extracted the results of interest for the present study, i.e. testing for upregulation in layer II/III excitatory neurons or layer V/VI cortico-cortical projection neurons. Separately for each of these two cell types, we defined upregulated genes as those with FDR-adjusted P<0.01 and a positive t-score (expressed higher in this cell type against the others) (Supplementary Tables 5, 6). We then created a union gene set that included 2622 genes upregulated in layer II/III excitatory neurons and/or layer V/VI cortico-cortical projection neurons. For each of these 2622 genes, we returned to our spatial transcriptomic data and extracted the layer*lobe interaction P value, and identified 56 genes whose layer*lobe interaction P values were significant at FDR<0.01, in addition to being upregulated in layer II/III excitatory neurons and/or layer V/VI cortico-cortical projection neurons (Supplementary Table 7).

### Gene-set association analysis with brain and behavioural variability

We used previously published SNP-wise summary statistics from a genome-wide association study in 30,810 adults from the UK Biobank population dataset, to test whether the 56 genes defined in the section above contain genetic variants that are associated with white matter connectivity between left-hemisphere core language network regions (33) (four connectivity metrics: pars opercularis – superior temporal cortex; pars opercularis – middle temporal cortex; pars triangularis – superior temporal cortex; pars triangularis – middle temporal cortex) (Supplementary Figure 8). Genetic variants were mapped to each of the 56 genes based on National Center for Biotechnology Information build 37.3 gene definitions as implemented in the MAGMA software (75), including 50 kb upstream and 50 kb downstream of each gene. Separately for each white matter connectivity metric we then used MAGMA to derive a single gene-based association P value per gene, using the default single-nucleotide polymorphism (SNP)-wise mean model. This analysis combined the association signals of all SNPs within a given gene, while considering the linkage disequilibrium between SNPs.

Separately for each of the four white matter connectivity metrics, the resulting gene-based association P values were used as input to test for a set-level association of the 56 genes defined in the previous section (above), using the GAUSS software (47). GAUSS has the advantage that it additionally identifies the subset of ‘driver genes’ with the maximal evidence of association, that best account for a significant set-level association. GAUSS uses a self-constrained null model for well-controlled type I error. We then adjusted the set-level association P values by Bonferroni correction for the four white matter connectivity metrics.

Finally, we ran the same procedure for the 56-gene set in relation to genetic association summary statistics from an analysis of word reading ability in 33,959 individuals carried out by the GenLang Consortium (34), and a study of dyslexia in 51,800 adults who self-reported having a diagnosis versus 1,087,070 controls (35), carried out using data from 23andMe, Inc. We then adjusted the set-level association P values by Bonferroni correction for these two language- and reading-related traits. (Adjusting the set-level association P values for six traits, i.e. four white matter connectivity metrics and two reading-related behavioural traits, made no difference to which associations were significant at adjusted P<0.05).

## Acknowledgements

This research was funded by the Max Planck Society (Germany) and the Netherlands Organization for Scientific Research (Language in Interaction consortium: Gravitation grant number 024.001.006). The funders had no role in study design, data collection and analysis, the decision to publish or preparation of the manuscript. We would like to thank the research participants and employees of 23andMe, Inc. for making this work possible.

## Data and Materials Availability

The spatial transcriptomic dataset created for this study can be found within the Gene Expression Omnibus with accession number <<TO BE ADDED>>. Genome-wide association summary statistics from the 23andMe study of dyslexia should be requested from 23andMe (https://research.23andme.com/dataset-access/). Other data sources are cited in the Methods section and can be accessed via the corresponding publications. This study used openly available software as cited in the Methods section. Additional code that supported the analyses in this study is available at <<LINK TO BE ADDED>>.

## Author Contributions

Conceptualization: SEF, WDJvdB, LEJ, CF. Data curation: MMKW, ZS, NV, SvH, LEJ. Formal analysis: MMKW, ZS. Funding acquisition: SEF, CF. Investigation: MMKW, ZS, LL, XZK, NV, SvH, LEJ. Methodology: MMKW, ZS, LL, XZK, NV, SvH, WDJvdB, LEJ. Project administration: CF. Resources: MM, WDJvdB, LEJ, SEF, CF. Supervision: SEF, CF. Visualization: MMKW, XZK, ZS, NV. Writing - original draft: MMKW, ZS, CF. Writing - review & editing: All authors.

## Supplementary Information

**Supplementary Figure 1.**
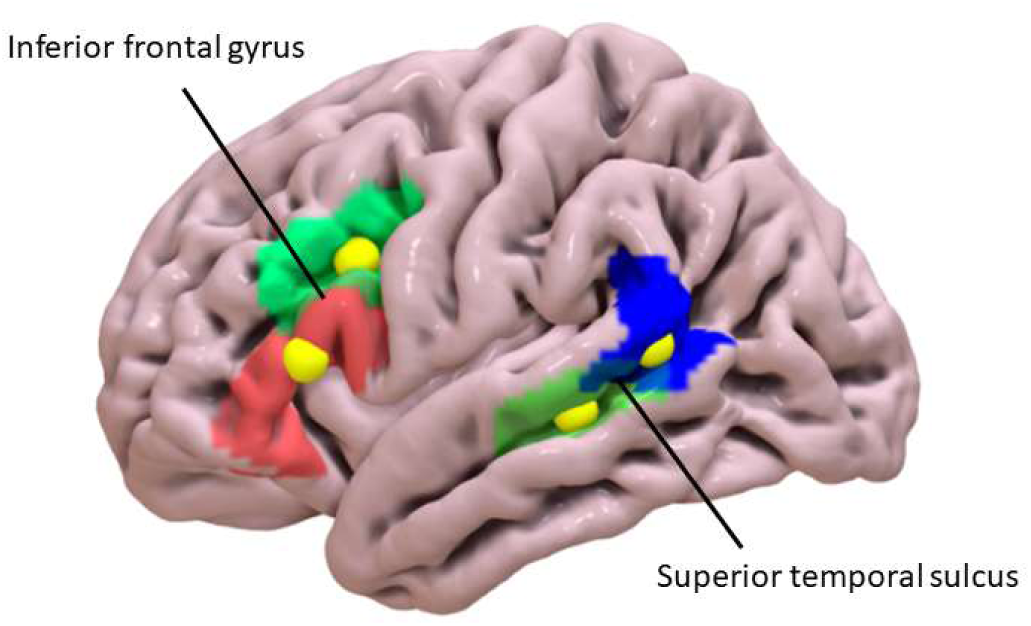
Schematic of the left-hemisphere regions sampled in this study. Two tissue blocks were taken from the inferior frontal gyrus. Blocks were approximately centred on the yellow spots indicated (block ‘gfi1’ more anterior/inferior, block ‘gfi3’ more posterior/superior). Two tissue blocks were also taken from the superior temporal sulcus. Again, blocks were approximately centred on the yellow spots indicated (block ‘gts4’ more anterior/inferior, block ‘gts5’ more posterior/superior). The broader coloured regions around the yellow spots represent four areas defined in the SENtence Supramodal Areas AtlaS (SENSAAS) (Labache et al. (2019) – see reference in the main manuscript).

**Supplementary Figure 2.**
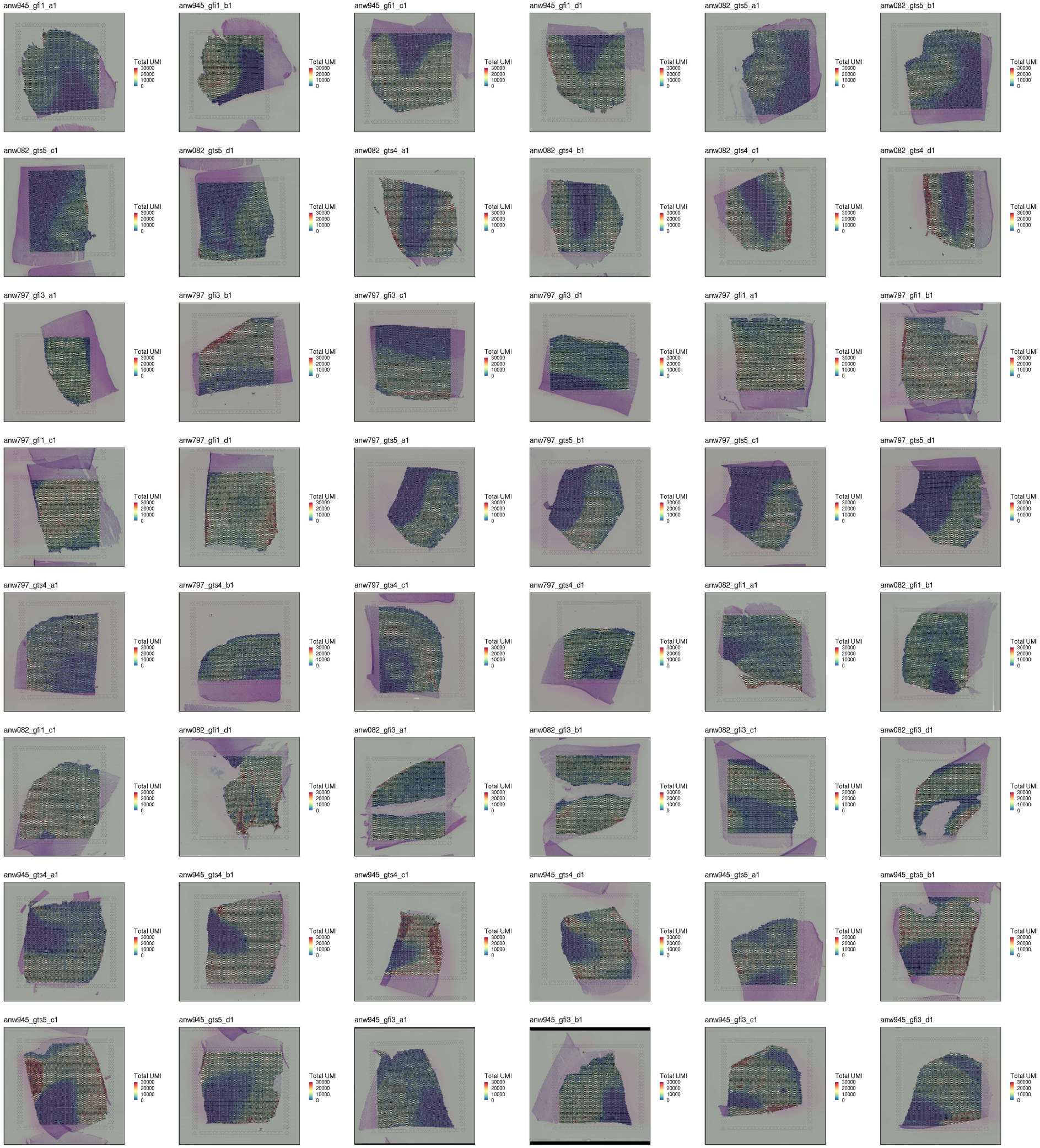
Unique molecular identifier counts across all spots for 48 cortical tissue sections. The pink/purple areas were outside of the capture area for spatial transcriptomics. The labels of the 48 cortical tissue sections are given as Donor_Block_Section, where the three donors were anw082, anw797 and anw945, the blocks were gfi1 & gf13 from the inferior frontal gyrus and gts4 & gts5 from the superior temporal sulcus, and the tissue sections from each block are called a1 & a2 (the first adjacent pair of sections) and a3 & a4 (second pair of adjacent sections).

**Supplementary Figure 3.**
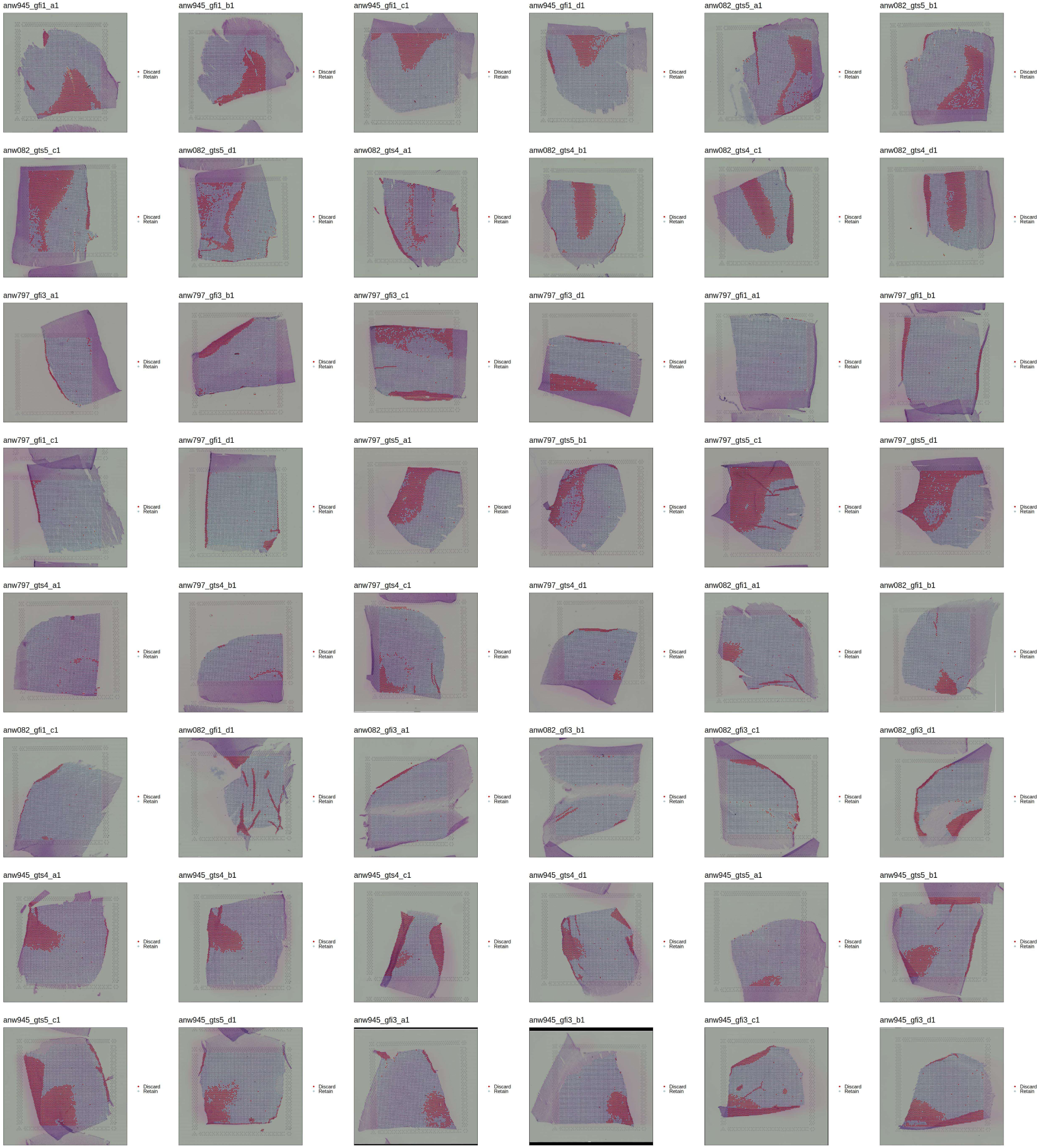
Spots excluded during spot-level quality control. Spots coloured red were excluded according to the process described in the main text (Methods). The labels of the 48 cortical tissue sections are given as Donor_Block_Section, where the three donors were anw082, anw797 and anw945, the blocks were gfi1 & gf13 from the inferior frontal gyrus and gts4 & gts5 from the superior temporal sulcus, and the tissue sections from each block are called a1 & a2 (the first adjacent pair of sections) and a3 & a4 (second pair of adjacent sections).

**Supplementary Figure 4.**
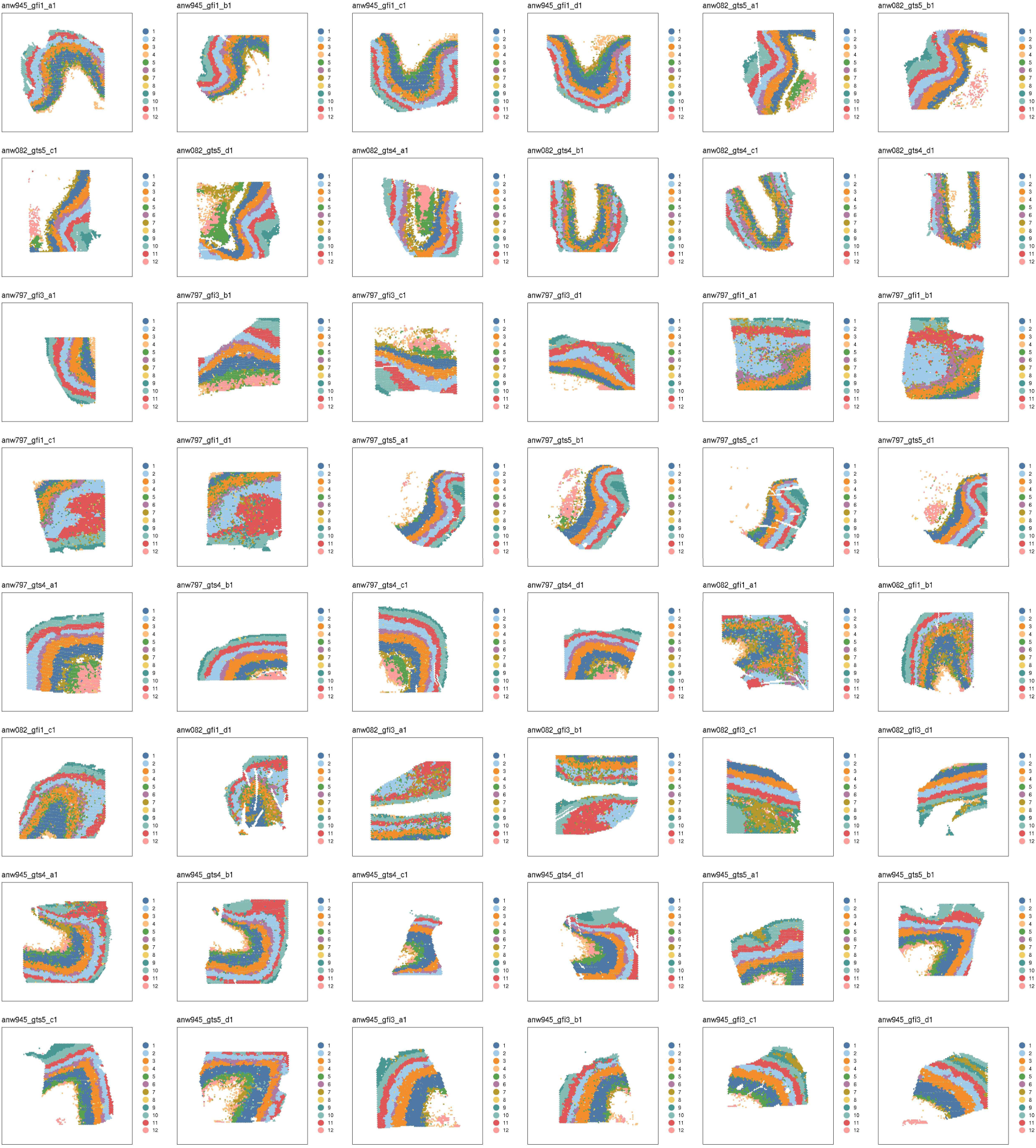
Spatial distributions of gene expression data-driven clusters of spots across 48 tissue sections. The labels of the 48 cortical tissue sections are given as Donor_Block_Section, where the three donors were anw082, anw797 and anw945, the blocks were gfi1 & gf13 from the inferior frontal gyrus and gts4 & gts5 from the superior temporal sulcus, and the tissue sections from each block are called a1 & a2 (the first adjacent pair of sections) and a3 & a4 (second pair of adjacent sections).

**Supplementary Figure 5.**
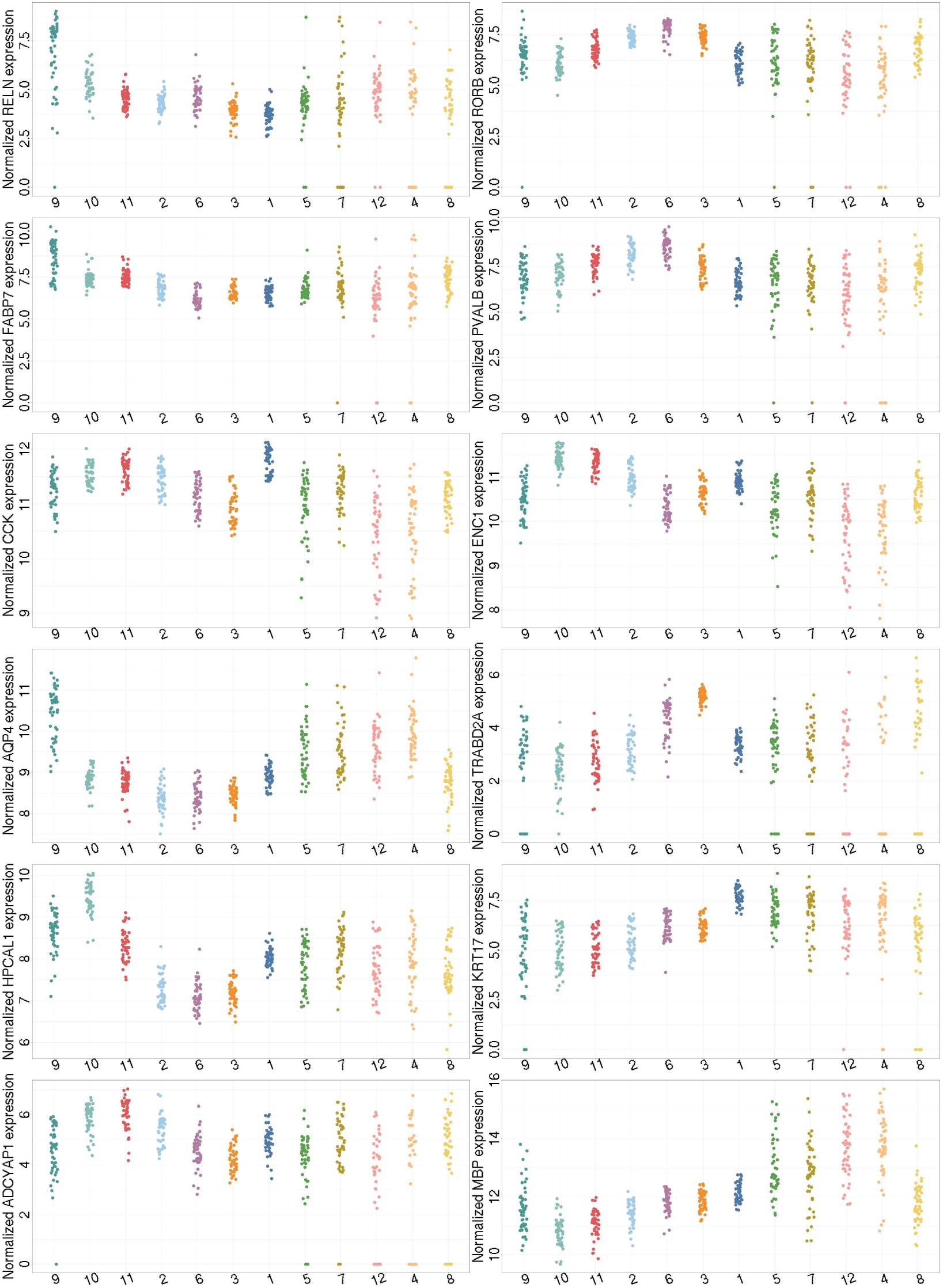
The expression levels of layer marker genes in data-driven clusters. Each panel shows the expression of a single marker gene, with the x-axis showing the data-driven clusters and the y-axis showing the normalized gene expression across 48 tissue sections. Clusters are ordered from left-to-right according to their laminar spatial locations from upper to lower. These data support the following correspondence: layerI=cluster9, layerII=cluster10, layerIII=clusters11&2, layerIV=cluster 6, layerV=cluster 3, layerVI=cluster1, with other clusters corresponding to white matter or with sporadic spatial distributions.

**Supplementary Figure 6.**
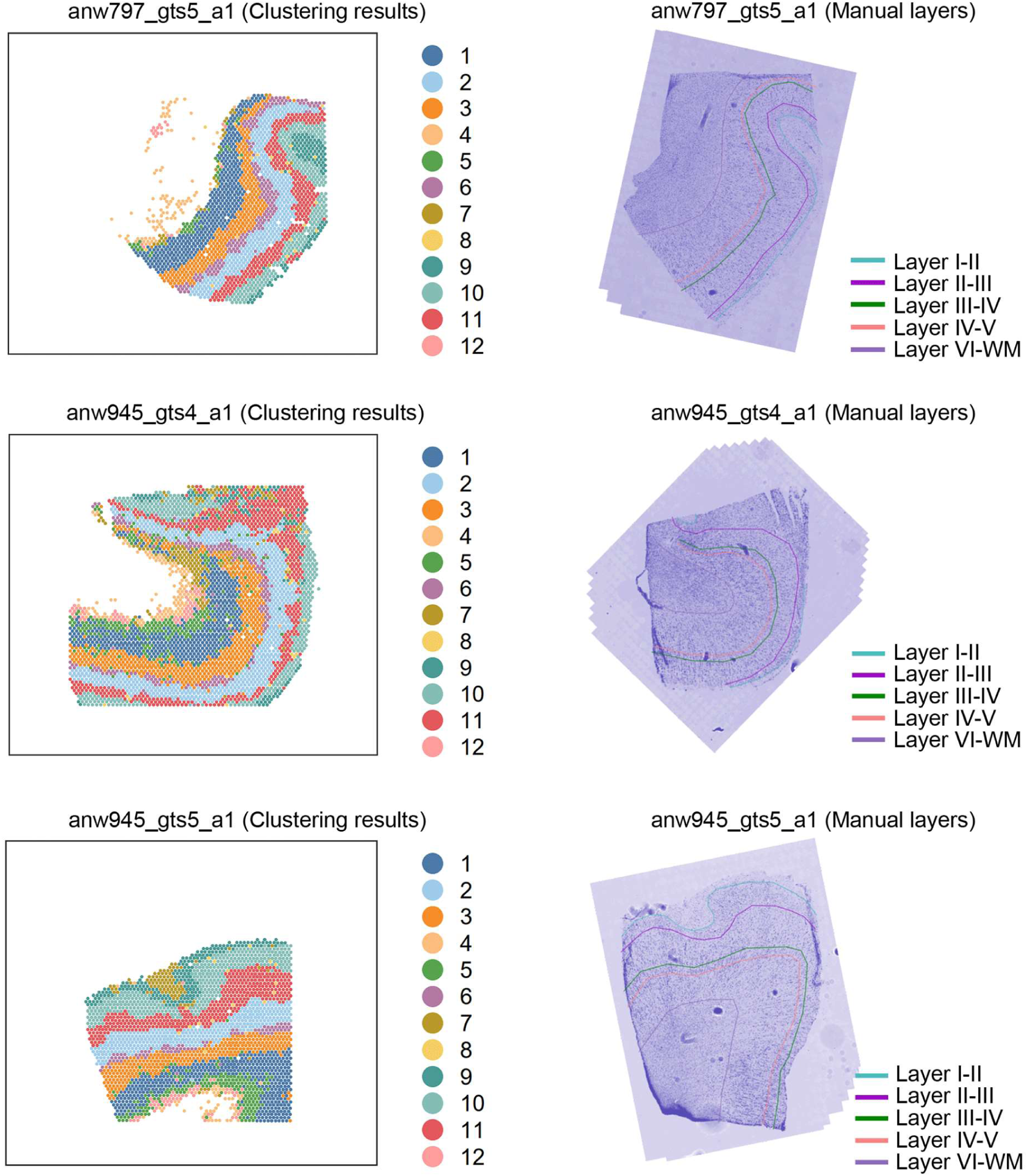
Cluster-layer correspondence assessed through cytoarchitecture. Comparison between data-driven gene expression clusters from the spatial transcriptomic data, and manually-defined cortical layers based on cytoarchitecture, across three pairs of tissue sections.

**Supplementary Figure 7.**
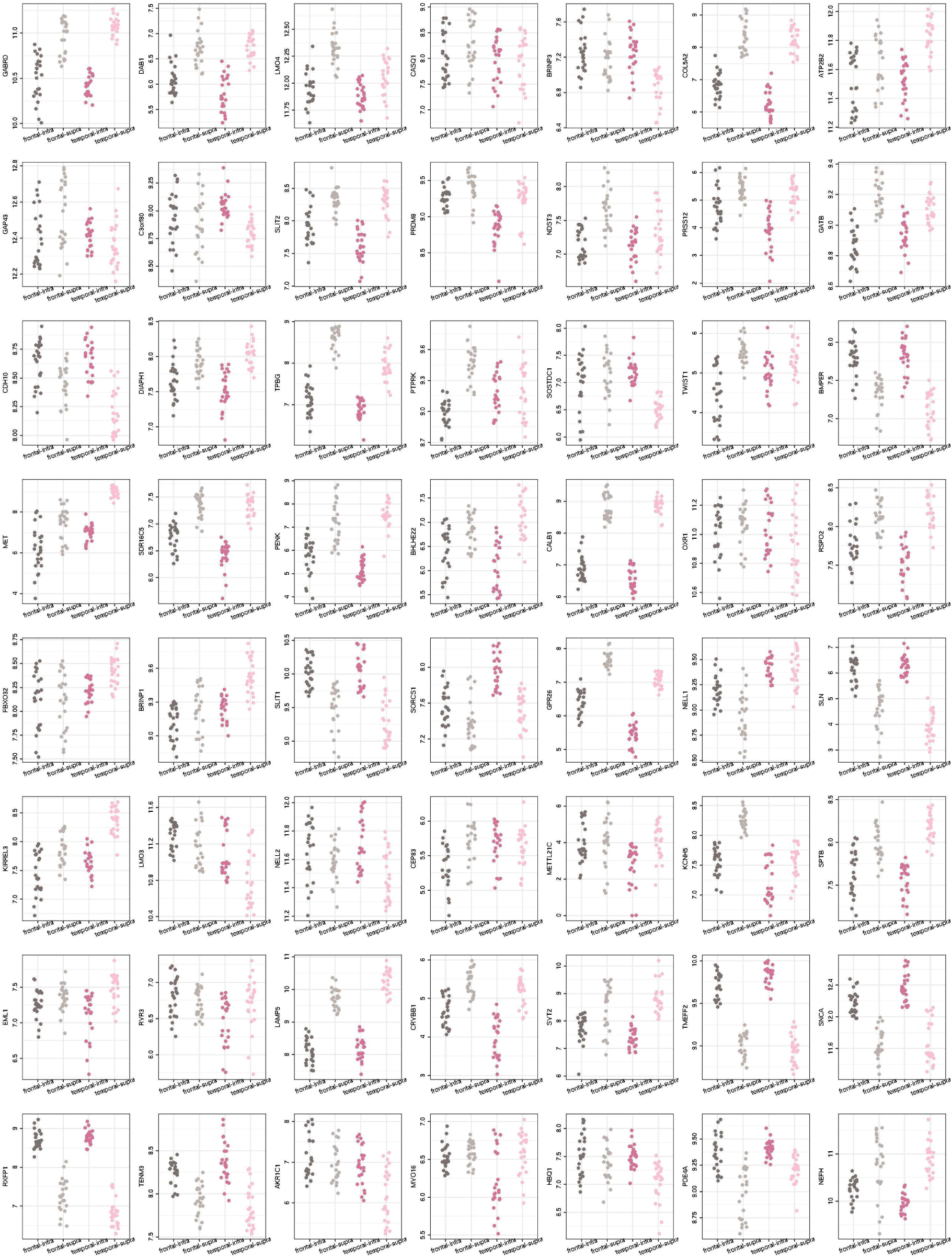
Significant layer*lobe interaction effects for 56 genes that also showed upregulation in Layer II/III excitatory neurons and/or Layer V/VI cortico-cortical projection neurons. Each panel shows the expression of a single gene, with the x-axis showing four pseudo-bulked clusters (see main text) and the y-axis showing the normalized gene expression across 48 tissue sections.

**Supplementary Figure 8.**
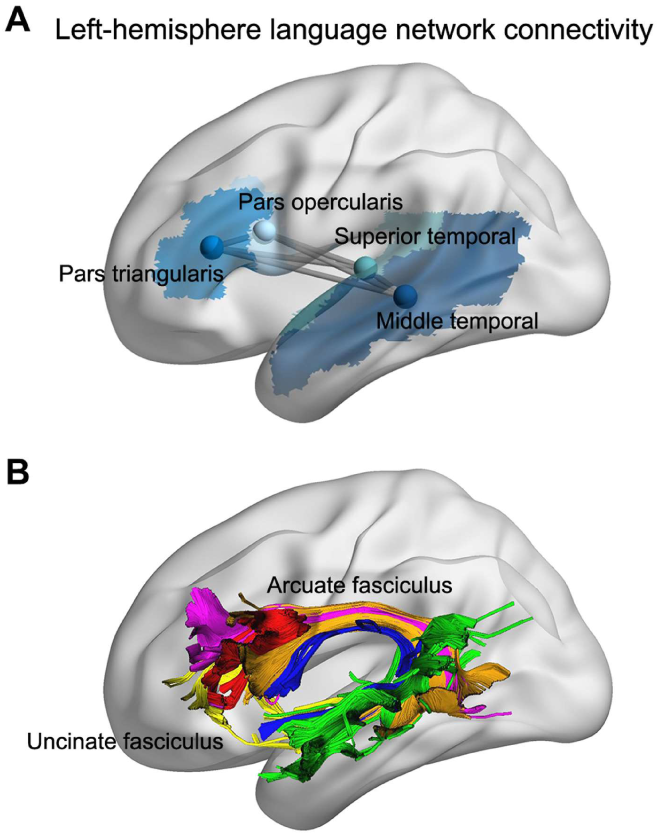
White matter connectivity between four regions of the core left-hemisphere language network. (A) The four cortical regions are shown in different shades of blue, as defined in the Automated Anatomical Labelling brain atlas (Tzourio-Mazoyer et al. 2002), and applied in the genome-wide association study of Sha et al. (2023) in 30,810 adults (see reference list in the main manuscript). Also shown is an abstract network representation where the four cortical regions are nodes and the structural connections between them are edges. In the present study we were interested in the four frontal-temporal network edges, i.e. pars opercularis - superior temporal cortex; pars triangularis - superior temporal cortex; pars opercularis - middle temporal cortex; and pars triangularis - middle temporal cortex. (B) Visualization of the white matter connections between the four cortical regions in an example individual, with gold representing connections between the pars opercularis and middle temporal cortex, blue representing connections between the pars opercularis and superior temporal cortex, purple representing connections between the pars triangularis and middle temporal cortex, and yellow representing connections between the pars triangularis and superior temporal cortex. Also shown are connections between the pars opercularis and pars triangularis (red), and connections between the middle temporal cortex and superior temporal cortex (green), but these within-lobe connections were not considered in the present study. Figure reproduced from Sha et al, (2023) under an open access Creative Commons Attribution License 4.0 (CC BY) (see reference list in the main manuscript).

